# ‘Are you even listening?’ - EEG-based decoding of absolute auditory attention to natural speech

**DOI:** 10.1101/2023.12.14.571397

**Authors:** Arnout Roebben, Nicolas Heintz, Simon Geirnaert, Tom Francart, Alexander Bertrand

## Abstract

**Objective:** In this study, we use electroencephalography (EEG) recordings to determine whether a subject is actively listening to a presented speech stimulus. More precisely, we aim to discriminate between an active listening condition, and a distractor condition where subjects focus on an unrelated distractor task while being exposed to a speech stimulus. We refer to this task as absolute auditory attention decoding.

**Approach:** We re-use an existing EEG dataset where the subjects watch a silent movie as a distractor condition, and introduce a new dataset with two distractor conditions (silently reading a text and performing arithmetic exercises). We focus on two EEG features, namely neural envelope tracking (NET) and spectral entropy (SE). Additionally, we investigate whether the detection of such an active listening condition can be combined with a selective auditory attention decoding task, where the goal is to decide to which of multiple competing speakers the subject is attending. The latter is a key task in so-called neuro-steered hearing devices that aim to suppress unattended audio, while preserving the attended speaker.

**Main results:** Contrary to a previous hypothesis of higher SE being related with actively listening rather than passively listening (without any distractors), we find significantly lower SE in the active listening condition compared to the distractor conditions. Nevertheless, the NET is consistently significantly higher when actively listening. Similarly, we show that the accuracy of a selective auditory attention decoding task improves when evaluating the accuracy only on the highest NET segments. However, the reverse is observed when evaluating the accuracy only on the lowest SE segments.

**Significance:** We conclude that the NET is more reliable for decoding absolute auditory attention as it is consistently higher when actively listening, whereas the relation of the SE between actively and passively listening seems to depend on the nature of the distractor.

## I. INTRODUCTION

The human auditory system is able to focus on a single speaker of interest while filtering out competing (auditory) stimuli, a process called selective (auditory) attention [1]. Recently, there has been an interest in algorithms that are able to decode this selective auditory attention based on neural activity, e.g. recorded with electroencephalography (EEG) [2]. In the literature, this task is often referred to as the ‘auditory attention decoding’ (AAD) problem [3]–[6]. However, we will refer to it as ‘selective’ auditory attention decoding (sAAD) to emphasize its goal of selecting one attended stimulus among multiple competing auditory stimuli. This terminology is introduced to distinguish it from another type of attention decoding, which we term ‘absolute’ auditory attention decoding (aAAD). In aAAD, the objective is to determine whether a subject is actively listening to *any* of the presented auditory stimuli or none of them [7]–[9]. Indeed, our brain is able to ignore all surrounding sounds, e.g., when focusing on an unrelated task, in which case an sAAD task would have no meaning. The sAAD problem, often referred to as just ‘AAD’ has been extensively studied, while the aAAD problem, which is a different variant of AAD, has received much less attention in the literature. Figures 1(a) and 1(b) illustrate this difference between the sAAD and aAAD problems.

**Fig. 1.**
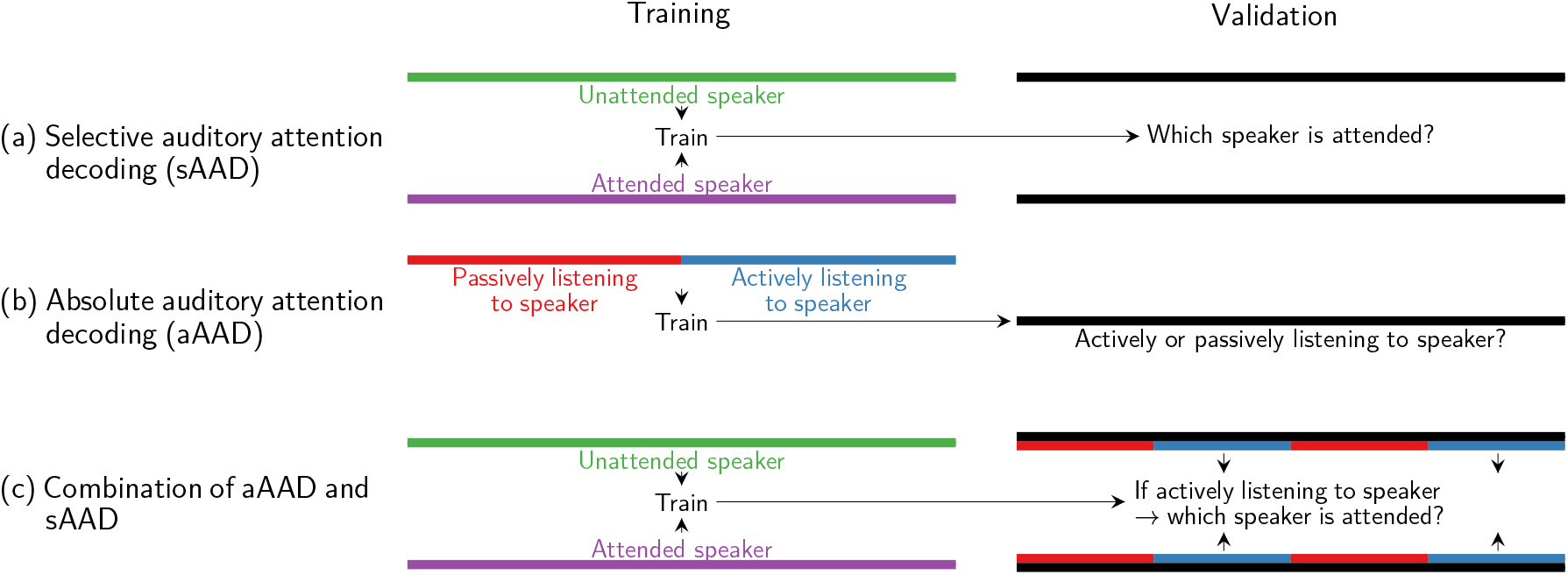
Overview of selective auditory attention decoding (sAAD), absolute auditory attention decoding (aAAD), and their combination. (a) In sAAD, multiple speakers are simultaneously present at each given moment. At training time, the audio from the attended and unattended speakers is leveraged, together with the neural activity of the subject listening, to train discriminative features and classifiers for sAAD. At validation time, these are subsequently used to discriminate which stimulus the subject is attending on held-out data. (b) Contrarily, in aAAD, the subject is either actively or passively listening to a speaker. At training time, the data are again leveraged to train the features and classifiers. At validation time, these are then used to assess whether the subject is actively listening or not on held-out data. (c) Both setups can be combined, where the sAAD accuracy is evaluated only on segments where actively listening is signalled by the aAAD algorithms, in order to avoid making arbitrary decisions due to the inattentiveness of the subject.

As shown in figure 1(a), more specifically, sAAD algorithms aim to discriminate between the attended and unattended speakers based on the subject’s neural activity and auditory stimuli of the speakers involved. At validation time, sAAD algorithms then aim to decode to whom the subject is listening. Contrarily, in the aAAD task (figure 1(b)), the algorithms are trained to detect the absolute state of auditory attention, i.e., to discriminate between an active or passive listening state of the subject. Note that while sAAD by definition assumes multiple competing speakers, aAAD can be performed in both a single– or a multi-speaker setting.

It has been shown that selective auditory attention can be decoded from EEG recordings, either based on the principle of neural envelope tracking (NET) [3], [10]–[12], or based on differences in the general neural activity [5], [6]. The former is based on the principle of neural response tracking. The neural response tracks certain characteristics of a speech stimulus, such as its envelope (i.e., the slow variations) over time [3], [9]–[12].

NET-based sAAD algorithms then exploit this neural tracking of the speech envelope being more strongly present for the attended speaker than for the unattended ones. As an alternative to NET, there exist sAAD algorithms that exploit differences in the neural activity itself, i.e., without using the auditory stimuli. For example, one can detect differences in the neural activity depending on the location of the attended stimulus [5], [6].

Literature about aAAD, i.e., discriminating actively versus passively listening is less extensive. In this work, we will extract two features from the EEG to perform aAAD, namely the NET and the spectral entropy (SE). As detailed supra, the NET characterises the tracking of the speech envelope by the neural response, and we hypothesize that this tracking disappears or decreases when the subject is not actively attending the speech. The SE, on the other hand, is a spectral EEG feature inspired by the concept of entropy, which is a measure of uncertainty of a random variable [13]. This SE has previously been hypothesised to predict regularity and predictability of the EEG [14]–[16].

Regarding the NET, Vanthornhout et al. [8] found significantly higher NET of subjects actively attending an auditory stimulus than that of subjects watching a silent movie, while ignoring the auditory stimulus. To quantify this NET, Vanthornhout et al. [8] trained linear decoders to reconstruct the speech envelope from the EEG. The resulting correlation between the reconstructed envelope and the ground truth envelope was shown to be significantly higher when actively focusing on the auditory stimulus. Kong et al. [7] investigated a similar scenario using the peak cross-correlation between the EEG and the speech envelope as a measure for the NET (without applying a neural decoder to potentially enhance this correlation). However, no difference across the active versus passive listening conditions was found in this study.

Regarding the SE, it has been shown experimentally that the SE was able to correctly predict anaesthetic depth [14], [15], respiration movements [15], sleep stages [15], and imagined finger movement [15], possibly due to changes in EEG regularity and predictability [14], [15]. Similarly, the SE allowed to discriminate subjects in rest from subjects performing mental arithmetic [17], and subjects in rest from subjects fixated on flashing patterns [16]. Lesenfants et al. [9] used the SE to quantify the level of sustained attention to an auditory stimulus. In this study, subjects were instructed to attend to an auditory stimulus, without any distractor conditions. The SE was then used to discriminate between high and low active listening segments, which are assumed to arise naturally as the subject’s focus might vary during the task. Lesenfants et al. [9] found that training linear NET decoders on high SE segments resulted in an improved NET decoding when evaluating on the full validation set. Given this improved NET decoding when training linear NET decoders on high SE segments, Lesenfants al. [9] concluded higher SE levels to correspond to higher auditory attention levels.

NET– and SE-based aAAD functionalities could be useful as an objective, general tool to measure the state of active listening during auditory EEG experiments. In addition, aAAD algorithms hold promise to be combined with sAAD algorithms. Indeed, advances in sAAD algorithms have recently been envisaged as a key ingredient towards, a so-called neuro-steered hearing device [2]. To improve speech intelligibility and speech quality, state-of-the-art hearing devices require speech enhancement algorithms to enhance the speaker of interest while suppressing other speakers. These speech enhancement algorithms currently use sub-optimal heuristics, such as assuming the user is facing the desired speaker, which are not always satisfied in everyday situations [2]. In neuro-steered hearing devices, the idea is to use sAAD algorithms to decode to whom the user is listening to control the speech enhancement algorithm of a hearing device, such that it can decide which speaker should be enhanced and which speakers should be suppressed in a multispeaker scenario [2]. Nevertheless, in these neuro-steered hearing devices, the sAAD algorithms consequently assume the subject is always actively listening to any one of the competing speakers. However, the sAAD algorithms should be disabled whenever the subject is not actively listening to any of the speech stimuli to avoid arbitrary speaker selections. Consequently, we will combine the aAAD and sAAD algorithms as illustrated in figure 1(c) by only evaluating the sAAD algorithm whenever active listening is signalled by the aAAD algorithm. In particular, we will investigate whether the NET and/or SE features are suited for aAAD in such a combined setting.

In summary, in this paper, we aim to decode whether or not a subject actively listens to a single speaker (aAAD) in different distractor conditions. To this end, we introduce a new EEG dataset with 10 subjects, wherein subjects are asked to either actively listen to a speech stimulus or to ignore it while silently reading a text or solving arithmetic exercises. Next to this dataset, we reuse a dataset from Vanthornhout et al. [8] in which the distractor condition consists of watching a silent movie. We then investigate whether both NET and SE can distinguish between the active listening condition and any of these distractor conditions. Finally, we combine this with an sAAD task with two competing speakers, in which the attended and unattended speaker needs to be discriminated. Segments labelled as passive listening by the NET and SE features are removed at validation time, attempting to take out segments in which the subject is not paying attention to any of the speakers, hence for which no truthful sAAD decision can be made.

Our research is complementary to, yet distinct from, the work of Vanthornhout et al. [8] and Lesenfants et al. [9]. First, we focus on natural speech within a broader scope of conditions, as opposed to the work of Lessenfants et al. [9], where no distractor conditions were present, and to the work of Vanthornhout et al. [8], where only a silent movie distractor condition was used and where only artificial, standardised sentences were used as speech stimuli at validation time, requiring little semantic processing in the brain. Second, we compare both the NET and SE features to discriminate between the active listening condition and distractor conditions, whereas the previous studies focused on either one of them and consequently do not have such a direct comparison. Finally, we apply aAAD to the sAAD task by only evaluating the sAAD accuracy on segments where the subject is signalled to listen actively, in order to support the use of aAAD in neuro-steered hearing devices.

We will demonstrate that higher SE does not necessarily correspond to higher auditory attention (as hypothesised by Lesenfants et al. [9]), i.e., SE shows different trends depending on the choice of the alternative (passive listening) condition. Contrary to the hypothesis of Lesenfants et al. [9] where actively listening was associated with a higher SE, we find a lower SE in subjects actively listening compared to subjects passively listening while watching a silent movie, performing arithmetic exercises and silently reading a text. Nevertheless, the sAAD accuracy decreases when only evaluating this sAAD accuracy on low SE segments. Consequently, the relation for SE between actively and passively listening seems to depend on the specific distractor task, rather than the absolute state of auditory attention. The SE, as such, seems to be a measure of cognitive resources spent on a specific task, rather than being a measure of auditory attention. The NET metric, on the other hand, shows consistent behaviour, i.e., it is always higher in the active listening condition compared to the passive listening condition, and the sAAD accuracy increases when only evaluating sAAD accuracy on high NET segments. NET therefore seems the better choice for aAAD.

This paper is structured as follows. First, we describe the algorithmic methodology to extract the NET and SE features for aAAD (Section II). Second, we briefly review the sAAD task and methodology (Section III). The datasets, data processing, and experimental setup are subsequently described (Section IV). The corresponding results are presented (Section V) and discussed (Section VI) thereafter. Finally, the conclusions are drawn (Section VII).

## II. ABSOLUTE AUDITORY ATTENTION DECODING

To decode whether a subject is in a state of active listening, i.e., perform aAAD, we will extract two features from the EEG:

1. **Neural envelope tracking (NET):** In the presence of a speech stimulus, the neural response tracks certain characteristics of that speech stimulus, such as its envelope, i.e., the slow variations over time [3], [9]–[12]. This envelope tracking is hypothesised to be more strongly present when the subject is more attentive to the speech [3], [8].
2. **Spectral entropy (SE):** Different tasks or conditions result in different neural activity across the brain [6], [9], [14], [16]–[18]. These differences in neural activity can, e.g., be quantified using the SE of the EEG recorded at a particular scalp location. The SE is hypothesised to quantify the predictability or regularity in the EEG activity, and quantifies the peakedness of the power spectral density (PSD) of the neural activity in the EEG indicating distinct oscillations, as will be detailed infra [9], [14]–[17]. Lesenfants et al. [9] hypothesised that the SE is higher in active listening compared to passive listening conditions.

The NET feature is discussed first (Section II-A), and the SE feature thereafter (Section II-B).

### A. Neural envelope tracking

To evaluate the degree of envelope tracking, a decoder consisting of a linear spatiotemporal filter is applied to the EEG signals to reconstruct the envelope of the speech stimulus. This is commonly achieved by minimising the squared error between the original and reconstructed speech envelope [3], [4], [8]. The degree of envelope tracking can subsequently be assessed by computing the correlation coefficient *ρ*(.) between the reconstructed and ground truth envelope.

Let *y*(*t*) represent the target speech envelope at sample time *t*, and *x*_*c*_(*t*) the sample of the *c*-th EEG channel at sample time *t*. Further, denote *C* as the number of EEG channels, *T* as the number of training samples, and *L* as the number of time lags. The goal is then to reconstruct the *T* subsequent target speech envelope samples **y** = [*y*(0) *y*(1) *… y*(*T −* 1)^⊤^ ∈ ℝ^*T*×1^] from channel-concatenated and timelagged EEG signals *X* ∈ ℝ^*T* ×*LC*^, defined as:

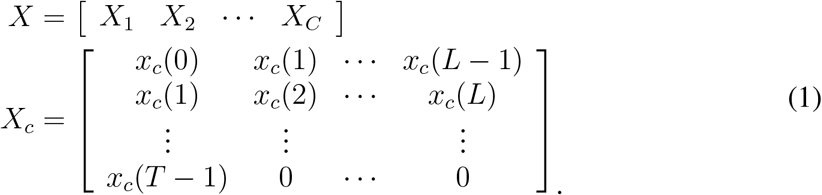

This reconstruction is achieved by designing a decoder 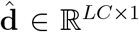 that minimises the squared error between the ground truth envelope **y** and the reconstructed envelope *X***d** [3], [19]:

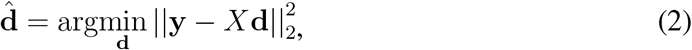

of which the solution equals [3], [19]:

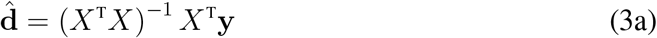

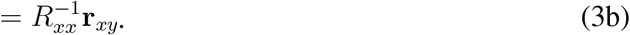

Herein, *R*_*xx*_ ∈ ℝ^*LC*×*LC*^ and **r**_*xy*_ ∈ ℝ^*LC*×1^ respectively denote the estimated EEG auto-correlation matrix and EEG-envelope cross-correlation vector.

At validation time, this decoder is applied to the validation data that were held out during training. The EEG data 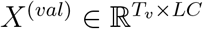 are then used to reconstruct the speech envelope 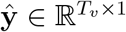 of length *T*_*v*_:

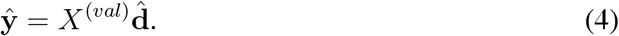

The correlation *ρ*(ŷ, **y**^(*val*)^) between this reconstructed envelope ŷ and the ground truth envelope 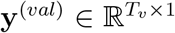, computed over *T*_*v*_ samples, is hypothesised to be higher when actively listening [8]. Therefore, we use this stimulus correlation as a first feature to discriminate between actively and passively listening. As per Vanthornhout et al. [8], the Spearman correlation coefficient, a rank-ordered Pearson correlation, is used in this work [20]. We will show that accordingly, the NET is higher when actively listening than when passively listening while performing arithmetic exercises and silently reading a text (Section V-A).

### B. Spectral entropy

The entropy is a measure for the uncertainty of a random variable by characterising the peakedness of the probability density function [13]. Herein, large entropy values map to high uncertainty and hence lower predictability. The entropy can also be applied to a normalised PSD^1^ in the frequency domain to describe the peakedness of this PSD [14], [17]. As the entropy is then applied in the frequency domain, it is referred to as spectral entropy (SE) [14], [17]. This so-called SE can be hypothesised to be used to leverage spectral differences, where higher SE levels correspond to less predictability in the neural activity. Let 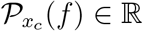 represent the normalised PSD estimate of the EEG signal *x*_*c*_(*t*) in channel *c*, then the SE in channel *c* and across the frequency band *f*_1_ − *f*_2_ is defined as:

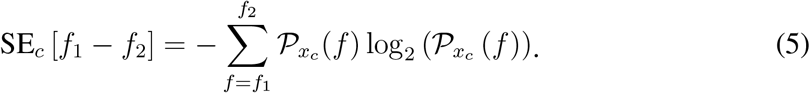

Previously, this SE allowed to the predict anaesthetic depth [14], [15], respiration movements [15], sleep stages [15], and imagined finger movement [15]. In addition, the SE allowed to discriminate between subjects in rest from subjects performing mental arithmetic [17], and subjects in rest from subjects fixated on flashing patterns [16]. It was hypothesised, that this predictive and discriminative power arises from the regularity and predictability differences in the EEG activity as characterised by the SE. Lesenfants et al. [9] hypothesised that higher SE values are associated with higher levels of auditory attention, as training decoders on the high SE segments outperformed the decoders trained on the low SE segments when evaluating on the full validation set. We will show that contrary to these results, lower SE is found when actively listening than when passively listening while watching a silent movie, silently reading a text and performing arithmetic exercises (Section V-A). Consequently, we will conclude that the SE seems to be a measure of cognitive resources spent on a specific task, rather than being a measure of absolute auditory attention.

III. APPLICATION TO SELECTIVE AUDITORY ATTENTION DECODING

As opposed to decoding when the subject is attentive or inattentive to a specific speech stimulus, in the sAAD setting, we aim to decode to which speaker the subject is attending in a multi-speaker scenario [2]–[4]. This corresponds to a selective auditory attention setup, as illustrated in figure 1(a). These sAAD algorithms are envisaged to be applied in so-called neuro-steered hearing devices, where the sAAD algorithms signal to the speech enhancement algorithm what speaker to enhance and what speakers to suppress [2].

As higher NET has been found for the attended speaker than for the unattended speakers, NET decoders can be used to discriminate between them [2]–[4]. The decoder for sAAD 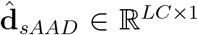 is computed using the envelope **y**_*a*_ ∈ ℝ^*T* ×1^ of the attended speaker, while the envelope(s) of the unattended speaker(s) is not used during training [2]–[4], [19]:

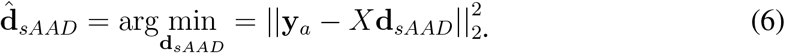

At validation time, the output of the decoder is correlated with the speech envelopes of all speakers over *T*_*v*_ samples. The speaker that yields the highest correlation *ρ*(.) between the reconstructed envelope 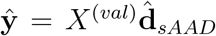 and the speaker envelope 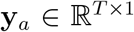 (for speaker *i*) is decoded as the attended one [3], [4], [19]. In the case of two speakers, the decision process can be described as:

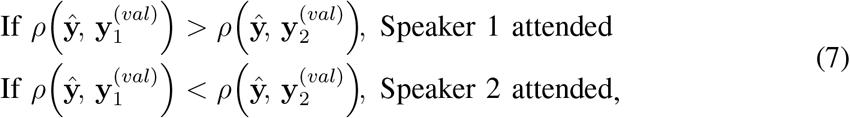

and the sAAD accuracy can be calculated as the percentage of correct decisions.

As illustrated in figure 1(c), both aAAD and sAAD could be combined in such a neurosteered hearing device. The aAAD algorithm could then be used to detect the segments where the subject is signalled to be in a state of active listening. By only using these segments corresponding to actively listening, we hypothesise that the sAAD decision process will improve the percentage of correct decisions.

## IV. EXPERIMENTAL PROCEDURES

To validate our methods, a new dataset was recorded, which is complementary to the datasets of Vanthornhout et al. [8] and Das et al. [21], which will also be used in our study. The dataset descriptions are given first (Section IV-A) and the description of how the corresponding data are processed thereafter (Section IV-B). The experimental setup is subsequently formulated (Section IV-C), as well as the statistical tests (Section IV-D).

### A. Datasets

In this work, we utilise three datasets: Dataset I is a newly recorded dataset that deals with subjects attending to an auditory stimulus versus subjects ignoring that stimulus while focusing on silently reading a text or solving arithmetic exercises. Dataset II originates from the work of Vanthornhout et al. [8] where subjects were either instructed to attend to an auditory stimulus or to ignore that stimulus while focusing on a silent movie. Finally, Dataset III is an sAAD dataset collected by Das et al. [21].

#### 1) Dataset I

##### Goal

The goal of this experiment is to investigate the neural differences between a setting where the subject is actively listening to a speech stimulus, and a distractor condition where the subject ignores the auditory stimulus while focusing on silently reading a text or solving arithmetic exercises.

##### Subjects

10 Dutch-speaking subjects (5 male, 5 female), between 21 and 27 years old, participated in the experiment. These subjects were unpaid volunteers. This study was performed in accordance with the Nurenberg Code. This human study was approved by Social-Societal Ethics Committee at KU Leuven – approval: G-2021-4407. All adult participants provided written informed consent to participate in this study.

##### Equipment

A 24-channel Smarting mobile EEG recording system was utilised [22]. The experiment was conducted in a non-radio frequency shielded room and the scalp of the subjects was treated with electroconductive gel while fitting the EEG cap. Raw EEG data were converted into MATLAB-compatible files using OpenVibe software [23].

##### Presentation structure

The speech stimuli consisted of children stories in Dutch [24], which were continuously presented to the subjects. While the speech stimuli were presented, the auditory attention of the subjects was manipulated by instructing the subjects to complete different tasks. To this end, the experiment consisted of three parts, possibly split into trials. During these parts and trials, the auditory attention was manipulated according to the following conditions, where the naming convention denotes to what task the subjects were instructed to pay attention to while the speech stimuli were continuously presented in all conditions:

- **‘Audio’**: The subjects were instructed to attend to the presented speech stimulus. To engage the subjects, they were told upfront that a questionnaire, concerning the content of that auditory fragment, had to be filled out at the end of the trial/part.
- **‘Mathematics’**: To reduce the auditory attention, the subjects needed to solve as many arithmetic exercises (e.g., 1012 *−* 448 = 564) as possible within one minute while ignoring the speech stimulus. The subjects were informed that no questions in the questionnaire would originate from those fragments of the story during which the arithmetic exercises had to be conducted.
- **‘Audio-Question’**: To increase the auditory attention, the subjects were instructed that a question was certainly to be asked about the content of this one minute fragment in the story.
- **‘Text’**: To reduce the auditory attention, the subjects were instructed to silently read a text while ignoring the speech stimulus. As before, subjects were informed in advance to fill in a questionnaire at the end of the part. In this condition, however, the questionnaire related solely to the content of the text, resulting in a no-attention condition to the speech.

An overview of the presentation structure of the conditions within each part and trial can be found in figure 2.

**Fig. 2.**
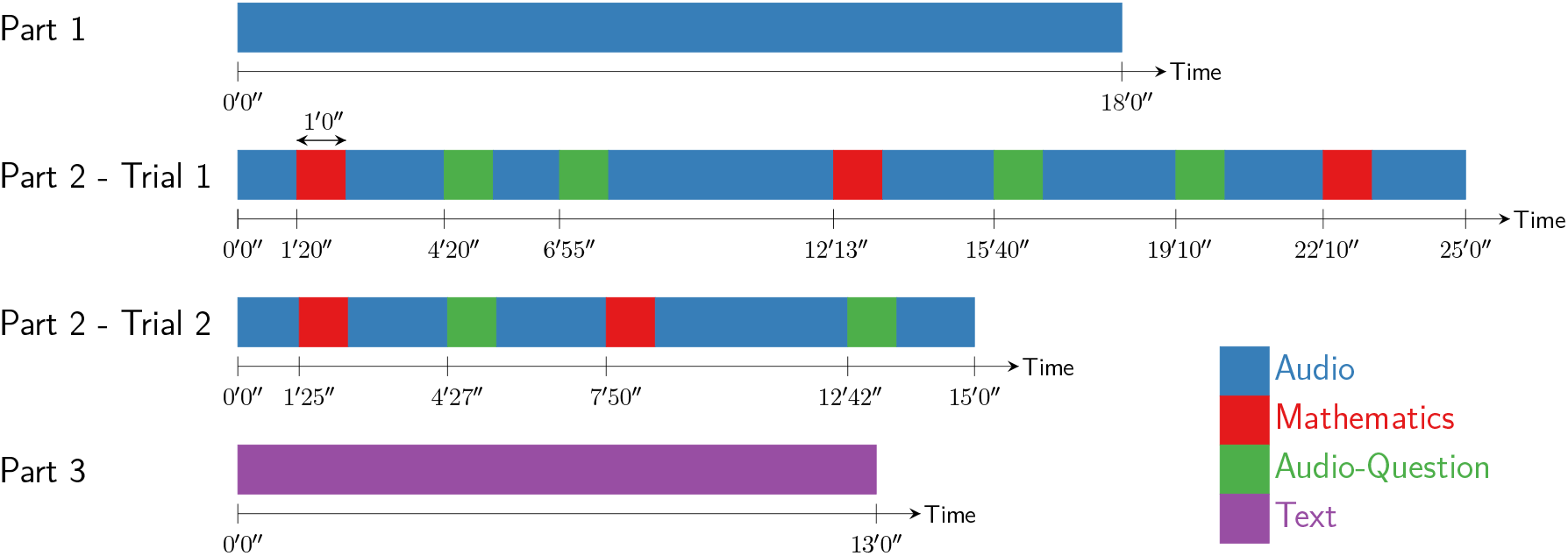
The presentation structure of Dataset I. The naming convention refers to the task the subjects were instructed to pay attention to while the speech stimuli were continuously presented in all conditions. During part 1, the subjects were instructed to attend an auditory stimulus and answer questions about the content afterwards (‘Audio’). During part 2, this structure was interleaved with specific tasks to be completed in order to modulate the subjects’ state of auditory attention. During the ‘Mathematics’ task, subjects needed to solve arithmetic exercises, reducing the auditory attention. During the ‘Audio-Question’ task, subjects needed to increase their auditory attention as a question certainly was to be asked about this fragment. Part 2 was split into two trials. During part 3, the subjects were instructed to silently read a text while ignoring the auditory stimulus.

The first part consisted solely of the ‘Audio’ condition, where the subjects were instructed to listen to the story ‘Bianca en Nero’ for 20 minutes. In the second part, the ‘Audio’ condition was interleaved with two different tasks, namely the ‘Audio-Question’ condition to increase the auditory attention and the ‘Mathematics’ condition to decrease the auditory attention. This part was split into two trials, corresponding to the lengths of the used speech stimuli, and to give the participants a break. The story ‘Ver van het kleine paradijs’ was used as speech stimulus in the first trial, and ‘Milan’ in the second trial. The final, third part consisted of the ‘Text’ condition, where the subjects were instructed to silently read a text while ignoring the speech stimulus. This text was provided as a printed document, and corresponded to a Dutch version of the short story ‘Vergif’ written by Roald Dahl [25]. The story ‘Eline’ was used as speech stimulus in this final part. Between each of the trials and parts, a short pause was inserted to give the subjects time to fill in the corresponding questionnaires.

The stories were played through the stereo speakers of a laptop. At the start of the experiment, the subjects could determine the volume themselves, such that they were comfortable during the experiments. The auditory stimuli were presented as is to the subjects without additional processing. The subjects were not familiar with the stories. The professional narrators were all male, except for the story used in the first trial of the second part. To keep the subjects motivated throughout the experiment, the subjects were informed that a gift card was to be handed out to the three best-scoring participants on the combined results of the questionnaires and arithmetic exercises.

#### 2) Dataset II

Dataset II is a subset of the dataset collected by Vanthornhout et al. [8]. This subset consists of 7 normal-hearing subjects, who all participated in two experiments. In the first experiment, subjects were instructed to attend a Dutch continuous auditory stimulus (story ‘Milan’). In the second experiment, the subjects were instructed to attend a silent, subtitled cartoon movie while ignoring an auditory stimulus. For each subject, about 15 minutes of data are available for both conditions. The EEG data consist of 64 channels. The interested reader can find a more detailed description of this dataset in the work of Vanthornhout et al. [8].

#### 3) Dataset III

Dataset III corresponds to the sAAD dataset as described in the work of Biesmans et al. [4], and as publicly available [21]. Herein, 16 normal-hearing subjects were exposed to a competing speaker scenario with two speakers, and were instructed to attend to one of the speakers while ignoring the other one. As auditory stimuli, four Dutch children’s stories were used. In total, approximately 72 minutes of recorded data per subject are available, using a 64-channel EEG cap. A more detailed description of this dataset can be found in the work of Biesmans et al. [4].

In the present study, we do not further consider the segments corresponding to the ‘Audio-Question’ condition of Dataset I (green segments in figure 2), to ensure a consistent active listening condition between Dataset I and Dataset II. Furthermore, the data of the text reading and arithmetic exercise solving distractor conditions are considered as one distractor condition, and hence their data are concatenated to increase the amount of data. We justify this approach since we want to binary discriminate between the active listening condition and the passive listening condition, although the amount of data for the text reading distractor condition is larger.

#### B) Data processing

##### 1) Preprocessing

Preprocessing is applied after the data collection to the EEG in order to remove artefacts, and to the speech in order to extract the envelope before computing the NET and SE features. The preprocessing framework is chosen the same as described in the work of Vanthornhout et al. [8], yet, with an additional muscle artefact removal step. The speech envelope is extracted from the audio data based on the procedure proposed by Biesmans et al. [4]. First, the raw audio data are filtered using a gammatone filterbank consisting of 28 filters, with centre frequencies between 50 Hz and 5 kHz, spaced according to 1 equivalent rectangular bandwidth [26], [27]. The output signal of each filter is thereafter transformed with a power law, i.e., |*y*_*k*_(*t*)|^0.6^, *k* = 1, …, 28. The signals are then linearly combined with equal weight and downsampled to 256 Hz, with the built-in low-pass anti-aliasing filter in MATLAB 2021b, to extract the envelope.

The EEG data are first also downsampled to 256 Hz. Next, muscle and eye artefacts are removed using a multichannel Wiener filter (MWF) approach as described by Somers et al. [28], [29]. Since the MWF is a data-driven filter, the data for all conditions for the same subject are filtered using the same MWF in order to avoid condition-dependent preprocessing. This approach requires artefact annotation, for which heuristic detection mechanisms are utilised [8], [30], further specified in A. The EEG data are subsequently re-referenced to the Cz-channel. Both the EEG and envelope are either bandpass filtered using a Chebyshev type 2 filter with cutoff frequencies tailored to the delta band (0.5−4 Hz) (for the NET calculation), or passed through unfiltered (for the SE calculation), as the SE frequency selection will be performed directly on the PSD in correspondence to (5). Finally, both signals are downsampled to 128 Hz and periods of longer silence (>0.25 s) are removed.

Additionally, for Dataset I, the EEG data are first linearly detrended to compensate for the strong baseline drift. The first downsampling before artefact removal is dropped for Dataset III, as the EEG data in this case are only available at 128 Hz and highpass filtered above 0.5 Hz [21].

##### 2) Hyperparameters

The following hyperparameters are adhered to:

- **NET:** The lag value *L* is chosen equal to *L* = 64, corresponding to a time window of 500 ms, to capture the relevant neural response [8], [12]. Similar to Vanthornhout et al. [8], all EEG channels are included in the design of the decoder. For the two 64-channel EEG datasets (Dataset II and Dataset III), the correlation matrix in (3b) is of dimension *LC* × *LC* where *LC* = 4096, which is quite large. For the sake of numerical stability and to avoid overfitting, we, therefore, apply L2-regularisation 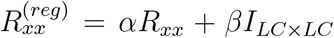, wherein *α, β* are computed according to the method presented by Ledoit and Wolf [31], of which a MATLAB implementation is available [32].
- **SE:** In correspondence to the work of Lesenfants et al. [9], the SE is calculated in a network of frontal, parietal-occipital and occipital channels (Fp1, Fpz, Fp2, AF7, AF3, AFz, AF4, AF8, F7, F5, F3, F1, Fz, F2, F4, F6, F8, PO7, PO3, POz, PO4, PO8, O1, Oz and O2 for the 64-channel EEG cap and Fp1, Fp2, F7, F8, Fz, O1 and O2 for the 24-channel EEG cap) on a frequency range spanning the alpha (8*−*13 Hz) and beta (13*−*30 Hz) bands, i.e., setting *f*_1_ = 8 Hz and *f*_2_ = 30 Hz in (5). The average of the SE values over these channels is utilised as an active listening feature. The PSD estimate is calculated using the multitaper spectral analysis using 7 Slepian tapers with a frequency spacing of 0.5 Hz [33].

#### C. Experimental setup

##### 1) Absolute auditory attention decoding

To study how the NET and SE features vary across the different active listening and distractor conditions, a 10-fold cross-validation is conducted per subject on Dataset I and Dataset II, where the folds are split chronologically. Regarding the NET, the decoder is trained solely on the active listening data of the left-in folds in order to reconstruct the speech envelope when attending to the speech stimulus, whereas regarding the SE, no training is required. The left-out fold is partitioned into windows, quantifying the amount of time given to decide whether the subject is in a state of active listening. To this end, window lengths of 5 s, 10 s, 20 s, 30 s and 60 s are used.

Subsequently, the classification accuracy of the features to discriminate between high versus low attention windows, defined as the ratio of the number of correctly predicted windows and the total amount of windows, is analysed. To this end, an equal amount of active and passive listening windows are present. A linear discriminant analysis (LDA) classifier is used, although other classifiers could be used as well [34]. This LDA classifier is trained using a nested 10-fold cross-validation procedure, where the folds are split chronologically. As for the NET, the covariance matrix estimate in the LDA classifier is regularised by adding a weighted identity matrix to the estimate according to the work of Ledoit and Wolf [31].

##### 2) Combination of aAAD and sAAD

Next, we aim to analyse whether aAAD can be incorporated into an sAAD framework, by evaluating whether the sAAD accuracy will increase when only making a decision when the subjects are identified to be most actively listening. A 10-fold cross-validation is conducted on the sAAD task of Dataset III, where the folds are split chronologically and the left-out folds are split into 60 s windows. The accuracy of these sAAD decoders is evaluated on a proportion of the left-out folds spanning the *x*%, *x* = 0, …, 100, of the segments signalled by the NET and SE features as having the highest absolute auditory attention. Regarding the NET, this selection of active listening segments is performed by applying the sAAD decoders on the left-out folds, computing the maximum of the correlation between the decoder output ŷ and both speech envelopes y_1_ and y_2_ 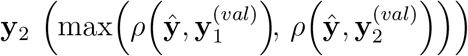 over 60 s windows, and selecting the segments which have the *x*% highest values of this feature. Regarding the SE, the *x*% lowest SE values on the left-out folds are selected in correspondence to the results on Dataset I and Dataset II as will be detailed infra. We hypothesise that the sAAD accuracy will subsequently increase as *x* decreases since no truthful sAAD decision can be made on segments where the subjects are not actively listening to any speech stimulus.

#### D. Statistical testing

Hypothesis tests are performed using the two-sided Wilcoxon signed rank test [35] and the resulting *p*-values are corrected for multiple comparisons using the Benjamini-Hochberg correction [36]. To assess the statistical significance of slopes, a linear regression model is fitted, and the t-test [37] on the coefficient corresponding to the slope is performed under the null hypothesis that the corresponding coefficient is zero. All hypothesis tests are performed with respect to a significance level *α* = 0.05.

The significance level for the classification accuracy is computed as the upper bound of a 95% one-sided confidence interval of a binomial distribution with success rate 50 % (chance level performance).

## V. RESULTS

### A) Absolute auditory attention decoding

Figure 3 shows the SE and NET correlations for each window of each subject (one data point = one window within a particular fold of one subject) and the per-subject averages across all windows in all folds on Dataset I and Dataset II for the 10 s and 30 s windows.

**Fig. 3.**
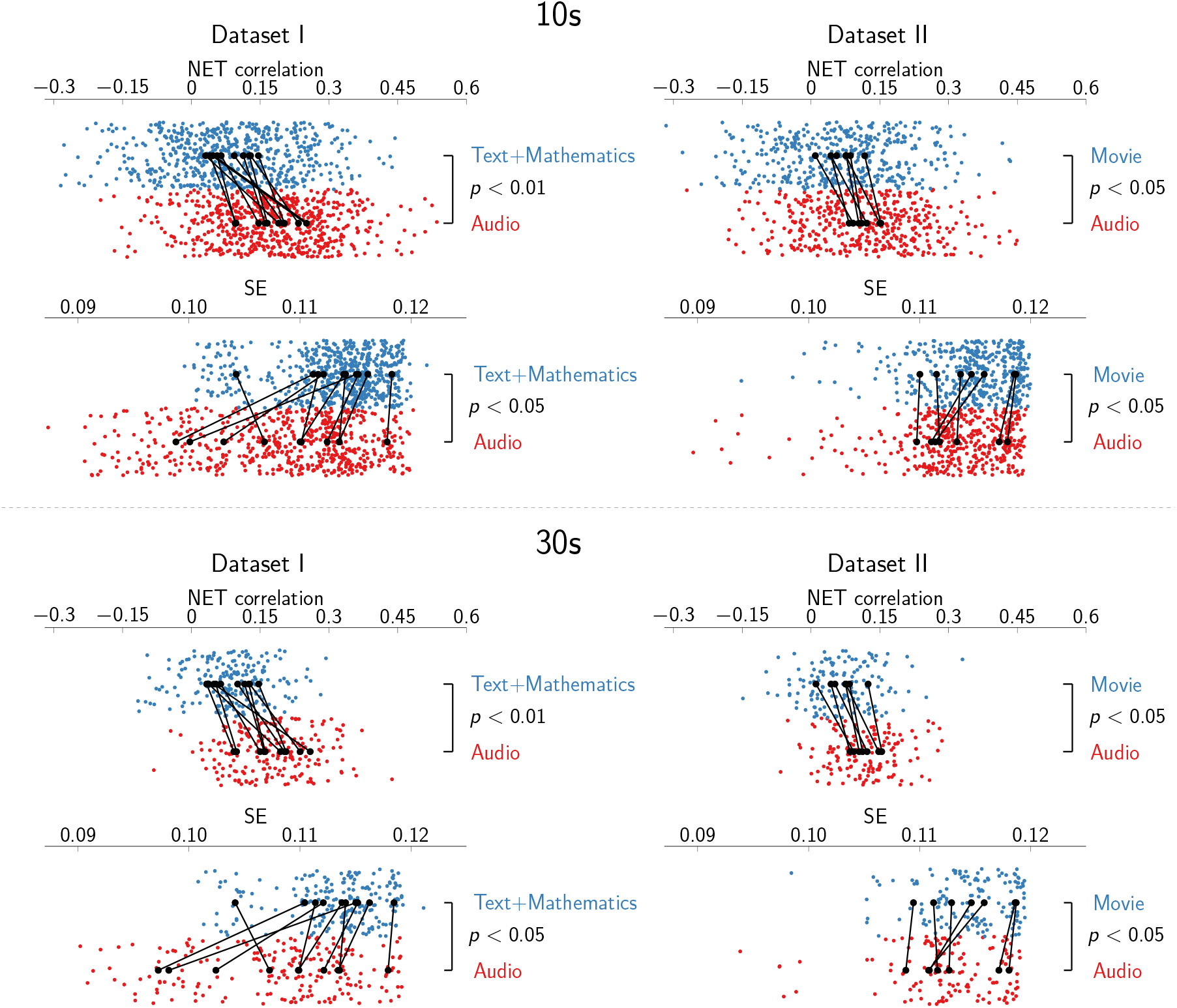
The neural envelope tracking (NET) correlations and spectral entropy (SE) on Dataset I and Dataset II. Contrary to the hypothesis of Lesenfants et al. [9], the SE is significantly lower when actively listening (‘Audio’) than when passively listening during the movie watching (‘Movie’), arithmetic exercise solving (‘Mathematics’) and text reading (‘Text’) distractor conditions. As hypothesised, the NET correlation is significantly higher in the active listening than in the distractor conditions, This experiment has been repeated for window lengths of 10 s and 30 s. Red and blue dots represent the individual data points for each combination of subject, fold, and window. The black dots represent the average feature value per subject after averaging across folds and windows. Lines connect these per-subject averages and *p*-values are noted on the right of the data.

The two-sided Wilcoxon signed rank test on the per-subject averages indicates a significant difference between the active listening and distractor conditions for the SE features (*p <* 0.05). However, the SE attains lower values in the active listening than in the distractor conditions, which is opposite from the hypothesis by Lesenfants et al. [9] where a higher SE was hypothesised for actively listening than passively listening (while not performing any distractor task).

Regarding the NET, significantly higher correlations are found for the active listening than for the distractor conditions (*p <* 0.05). These results are consistent with the work of Vanthornhout et al. [8], where higher NET correlations were found for subjects actively listening than for subjects passively listening while watching a silent movie. Nevertheless, while Vanthornhout et al. [8] used artificial, standardised sentences, we used natural speech and introduced two new distractor tasks.

When applying an LDA classifier on these features, we obtain classification accuracies as shown in figure 4. Both the NET and the SE features outperform chance level, and are subsequently able to discriminate between the active and passive listening condition, wherein the NET visually seems to suffer more from shorter window lengths than the SE.

**Fig. 4.**
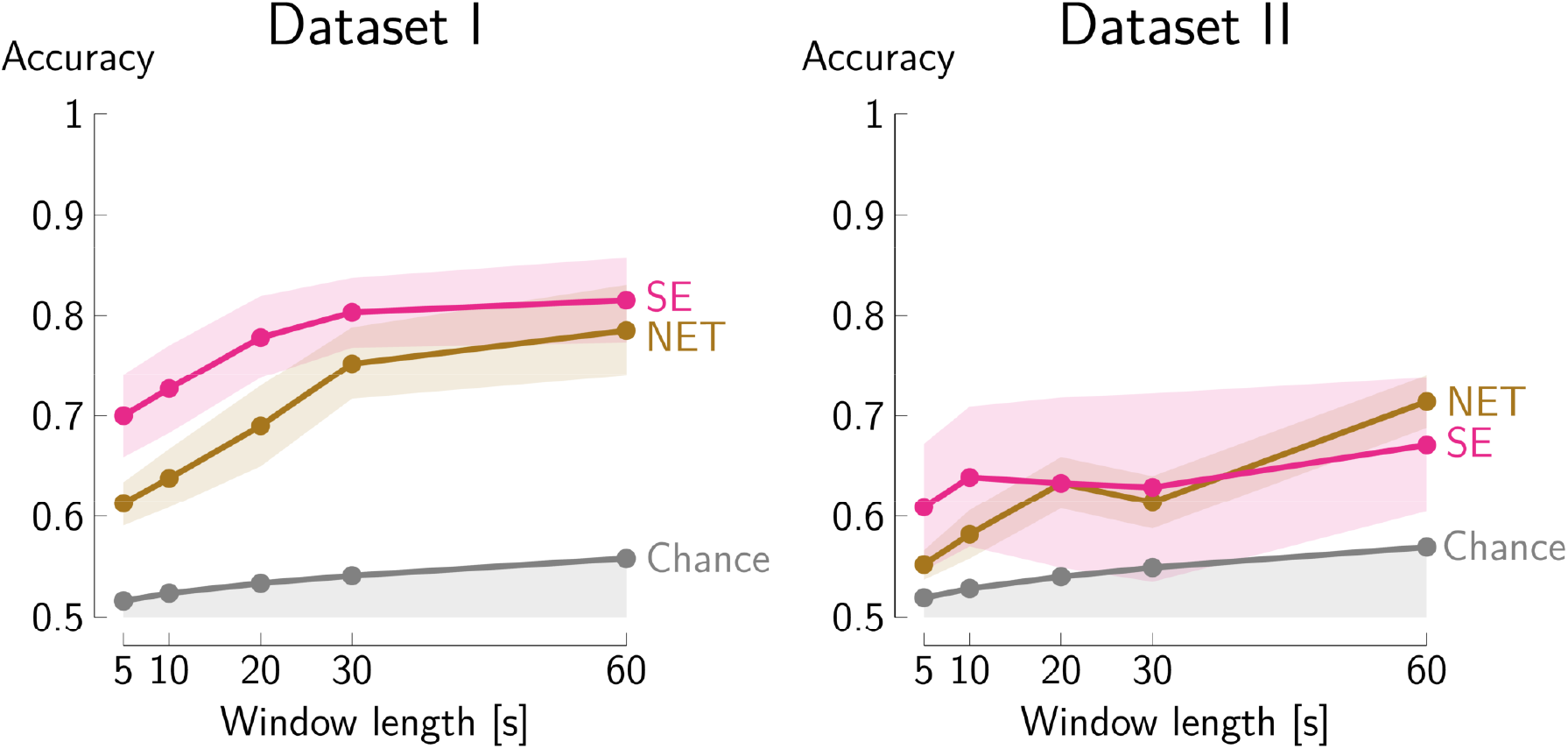
Classification accuracies across subjects, after averaging across folds, of the absolute auditory attention decoding (aAAD) task on Dataset I and Dataset II using the neural envelope tracking (NET) correlations and the spectral entropy (SE). The mean accuracy across the subjects is marked in bold and the standard deviation is shown as shading. Chance level is computed as the upper bound of a 95% one-sided confidence interval of a binomial distribution with success rate 0.5, and shown in shading, with the upper limit marked in bold. On both datasets, the methods yield accuracies above chance level.

In Dataset I, the standard deviation across subjects of the SE classification accuracy (0.14 (10 s), 0.11 (30 s)) is comparable to the NET classification accuracy (0.09 (10 s), 0.11 (30 s)). In Dataset II, however, this standard deviation across subjects of the SE classification accuracy (0.18 (10 s), 0.25 (30 s)) is larger than the standard deviation of the NET classification accuracy (0.06 (10 s), 0.07 (30 s)).

### B. Combination of aAAD and sAAD

Figure 5 shows the sAAD accuracy on Dataset III, when evaluated on the *x*%, *x* = 0, …, 100, segments with the highest absolute auditory attention, i.e., the *x*% segments with the highest NET correlation or the lowest SE, in correspondence with the results on Dataset I and Dataset II.

**Fig. 5.**
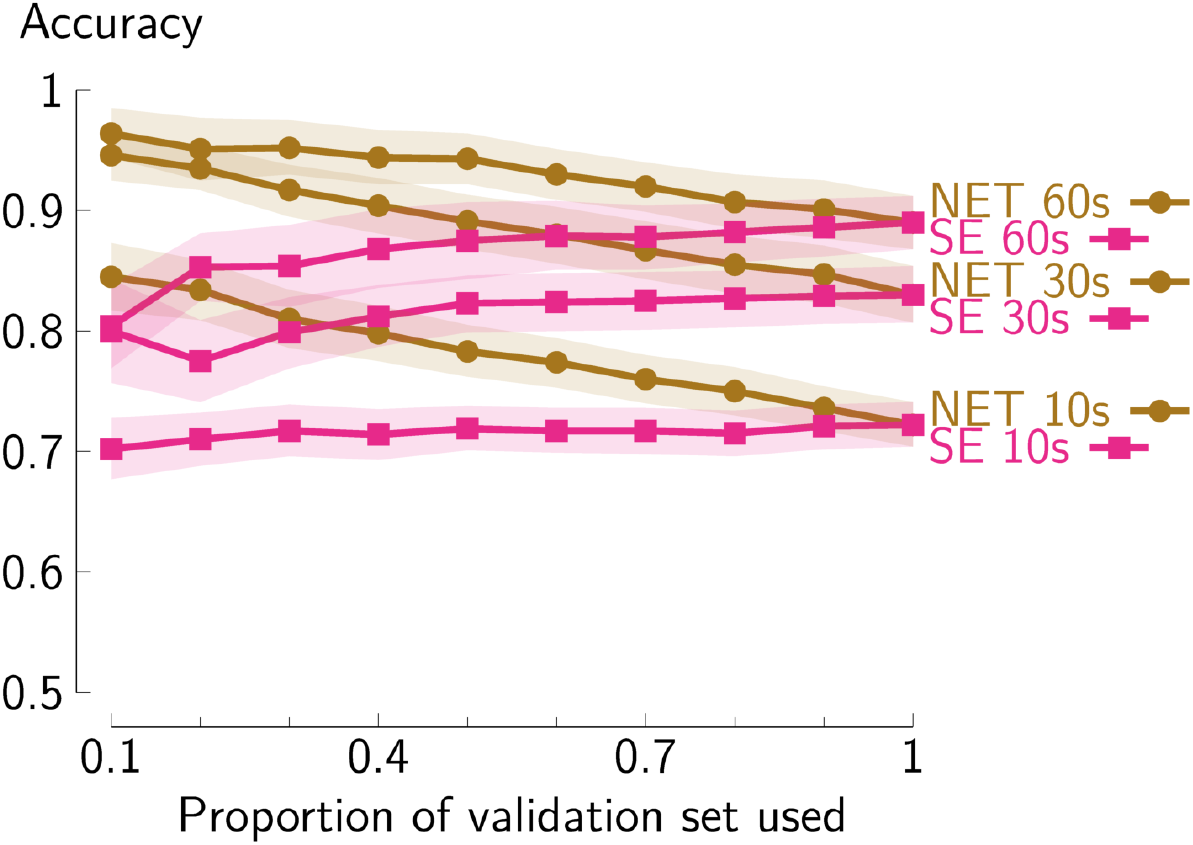
Average selective auditory attention decoding (sAAD) accuracies across subjects after averaging across folds. The sAAD accuracies are calculated on the *x*%, *x* = 0, …, 100, highest active listening segments of Dataset III, as signalled by the highest neural envelope tracking (NET) correlation and lowest spectral entropy (SE) segments conform the findings on Dataset I and Dataset II. Contrary to the results on Dataset I and Dataset II, a negative trend is observed when only evaluating the sAAD segments on a decreasing number of active listening segments as signalled by the lowest SE. By selecting the highest NET correlation segments, the sAAD accuracy increases with a decreasing proportion of the validation set used as hypothesised.

Contrary to the hypothesis, evaluating sAAD accuracy on the lowest SE segments shows a significant decrease in accuracy when decreasing *x* (*p <* 0.01). This trend is different than what is expected from the SE results on Dataset I and Dataset II, where lower SE was associated with more actively listening. This result is, however, consistent with the study of Lesenfants et al. [9], where a higher SE was hypothesised to correlate with higher auditory attention.

Nevertheless, evaluating the sAAD accuracy on the highest NET correlation segments, the sAAD accuracy increases as *x* decreases, illustrating the viability of this approach. This trend is, furthermore, significant (*p* < 10^−6^). This is consistent with the NET results on Dataset I and Dataset II, where a higher NET was observed in segments where the subject is actively listening to the speech.

## VI. DISCUSSION

Lesenfants et al. [9] hypothesised *higher* SE to be associated with *active listening* rather than with passive listening (not performing any interfering task) conditions. We find contrasting results: On the one hand, we find *lower* SE when *actively listening* versus passively listening while silently reading a text, performing arithmetic exercises (Dataset I), or watching a silent movie (Dataset II). On the other hand, we do find lower sAAD accuracies (Dataset III) when evaluating sAAD accuracy only on the lowest SE segments, which, by the findings in Dataset I and Dataset II, should correspond to segments of higher absolute auditory attention. Keeping only the lowest SE segment in Dataset III thus possibly keeps (rather than removes) segments of lower absolute auditory attention. In the sAAD setting where there is no dedicated distractor task, our results are actually consistent with the observations in the work of Lesenfants et al. [9], which also did not include a specific distractor condition.

Consequently, the relation of the SE between the active and passive listening conditions is inconsistent, and seems to depend on the distractor task. Our hypothesis is that the SE does not decode the absolute auditory attention, but rather reflects the differences in overall cognitive resources spent on each task (either active listening or a distractor condition). Indeed, in Dataset I and Dataset II, the passive listening conditions are all coupled to specific distractor conditions (movie watching, arithmetic exercise solving and text reading) that require mental resources. On the contrary, in Dataset III and the work of Lesenfants et al. [9], there is no specific distractor condition, such that within Dataset III and the work of Lesenfants et al. [9] a higher SE might be related to more actively listening and lower SE, e.g., to mind-wandering. The relation for SE between actively and passively listening, consequently, seems to depend on the specific distractor task. The SE, furthermore, cannot discriminate between different tasks giving rise to the same overall peakedness in the PSD. The practical relevance of the SE as a generic feature for absolute auditory attention decoding is therefore questionable. Nevertheless, the SE does prove useful in discriminating between two distinct, predefined tasks (e.g., actively listening versus watching a movie) whenever both tasks require a different amount of cognitive resources. However, the relative change of the SE between two conditions will depend on the nature of the task(s) in both conditions, and is thus not necessarily (directly) related to the state of active versus passive listening. This hypothesis is, furthermore, consistent with the predictive power of the SE in dedicated tasks, such as for anaesthetic depth [14], [15], respiration movements [15], sleep stages [15], and imagined finger movement [15], and consistent with the discriminative power in dedicated tasks, such as between subjects in rest and performing mental arithmetic [17], and between subjects in rest and fixated on flashing patterns [16]. Additionally, these findings further support the increasing awareness of the fact that general EEG features or models often do not generalise well across datasets and conditions, and can be heavily tailored to specific experimental conditions., e.g., [2], [38]–[40]

The NET correlations, on the other hand, do not suffer from this phenomenon of relative change between conditions since this technique directly measures the coupling between the acoustic stimulus and the neural response to that stimulus. Due to this explicit coupling, this trend is expected to be consistent across different experimental settings. We indeed observe that the NET correlations are consistently higher in the active listening condition compared to the passive listening conditions, for any of the distractor tasks, in line with the findings of Vanthornhout et al. [8]. Furthermore, the sAAD accuracy increases when evaluating sAAD on a reduced portion of the validation set as signaled by the highest NET correlations, keeping only high auditory attention segments. As a result, even though the NET-based classification does not outperform the SE-based classification in figure 4, the consistency of the NET feature across different settings makes it the better candidate for deployment in aAAD and sAAD tasks. Furthermore, it is noted that this NET accuracy can likely be improved by also including an encoder to the speech envelope next to the decoder on the EEG, resulting in a canonical correlation analysis (CCA) approach [41], [42].

These experiments, additionally, seem to provide a proof of concept for aAAD in general, and for using it in combination with sAAD, e.g., in the context of neuro-steered hearing devices. Future work could focus on validating these results on a dataset fully tailored towards combining sAAD and aAAD, where the auditory attention of subjects is manipulated with specific tasks during an sAAD experiment. To date, such a dataset does not exist to the best of our knowledge. Additionally, although the experiment of Dataset I was conducted in a non-radio frequency shielded classroom, both features significantly discriminate between the active and passive listening conditions. This indicates that this technology is viable in more realistic environments, outside controlled radio frequency shielded laboratories.

In summary, the NET feature seems better suited for aAAD and sAAD due to the consistent relation between actively and passively listening, independent of the type of passive listening condition. While the SE seems to be able to discriminate between two specific conditions, it does not seem useful as a general feature for aAAD due to the inconsistent relation between actively and passively listening across different conditions.

## VII. CONCLUSION

In this paper, we have discriminated between subjects actively listening and passively listening, a task referred to as absolute auditory attention decoding (aAAD). Both neural envelope tracking (NET) and spectral entropy (SE) features were used to perform this aAAD task. To this end, we have introduced a new dataset containing an active listening condition, as well as distractor conditions during which the subject silently reads a text or solves arithmetic exercises while passively listening to a speech stimulus. Next to this new dataset, we have also used an existing dataset where the distractor condition consisted of watching a silent movie. Contrary to previous work, where SE was hypothesised to be higher in subjects actively listening than in subjects remaining passive (without any distractor), we have found that the alpha and beta band SE in the frontal, parietal-occipital, and occipital channels increases from the active listening condition to the passive listening condition with distractor tasks such as movie watching, silently reading a text, and performing arithmetic exercises distractor conditions. On the contrary, the NET was found to be consistently higher between the active listening and any of the distractor (i.e., passive listening) conditions. Further, only evaluating selective auditory attention decoding (sAAD) accuracy on segments of high NET shows an increased sAAD accuracy, whereas evaluating on segments of low SE shows the reverse trend, thus possibly removing auditory attentive segments. Thus, likely, the SE rather relates to the cognitive load required for each task than the actual absolute auditory attention, whereas the NET correlations directly relate to (in)attention to the stimulus and are consistent across several datasets and tasks. The NET, thus, seems the better option to estimate absolute auditory attention, and appears to be more suited to use in conjunction with an sAAD task, e.g., in the context of neuro-steered hearing devices.

## APPENDIX

Segments are annotated as eye artefacts if the total power in the frontal channels (Fp1, AF7, AF3, Fpz, Fp2, AF8, AF4 and Afz for the 64-channel EEG cap, and Fp1, Fp2 and Fpz for the 24-channel EEG cap) is higher than 5 times the baseline power in those channels, according to the procedure of Vanthornhout et al. [8].

Regarding the muscle artefact detection, the EEG signal is first filtered using a zerophase Chesbyshev type 2 filter bandpass filter with cutoff frequencies between 20 Hz and 60 Hz. Segments are thereafter annotated as muscle artefacts if the total power of this filtered EEG signal in the channels at the side of the head (AF7, F7, F5, FT7, FC5, T7, C5, TP7, CP5, P7, P5, P9, PO7, AF8, F6, F8, FC6, FT8, C6, T8, CP6, TP8, P6, P8, PO8 and P10 for the 64-channel EEG cap, and F7, T7, CP5, F8, T8, TP9 and TP10 for the 24-channel EEG cap) is higher than 60 times the baseline power in those channels [30].

## ACKNOWLEDGEMENTS

We thank Ir. Linsey Dewit-Vanhaelen and Ir. Elly Brouckmans for writing the protocol and performing the experimental recordings for the new Dataset I. We thank Dr. Jonas Vanthornhout, Dr. Lien Decruy and Prof. Tom Francart for granting us access to Dataset II [8].

This research is funded by Aspirant Grant 1S31522N (for N. Heintz) from the Research Foundation – Flanders (FWO), a PDM mandate from KU Leuven (for S. Geirnaert, No. PDMT1/22/009), a junior postdoctoral fellowship fundamental research from the FWO (for S. Geirnaert, No. 1242524N), FWO project nr. G081722N, Internal Funds KU Leuven IDN project IDN/23/006, the European Research Council (ERC) under the European Union’s Horizon 2020 research and innovation programme (grant agreement No 802895 and grant agreement No 101138304), and the Flemish Government (AI Research Program). The scientific responsibility is assumed by its authors. Views and opinions expressed are those of the author(s) only and do not necessarily reflect those of the European Union or ERC. Neither the European Union nor the granting authority can be held responsible for them.

## Footnotes

1 The normalisation here refers to a division with the area under the curve of the PSD, such that its sum across the frequency axis is equal to one, similar to a probability density function.

## REFERENCES

[1] Cohen RA. Introduction. In: The Neuropsychology of Attention. Boston, MA: Springer US; 2014. p. 3–10.

[2] Geirnaert S, Vandecappelle S, Alickovic E, de Cheveigne A, Lalor E, Meyer BT, et al. Electroencephalography-Based Auditory Attention Decoding: Toward Neurosteered Hearing Devices. IEEE Signal Processing Magazine. 2021 Jul;38(4):89–102.

[3] O’Sullivan JA, Power AJ, Mesgarani N, Rajaram S, Foxe JJ, Shinn-Cunningham BG, et al. Attentional Selection in a Cocktail Party Environment Can Be Decoded from Single-Trial EEG. Cerebral Cortex. 2015 Jul;25(7):1697–706.

[4] Biesmans W, Das N, Francart T, Bertrand A. Auditory-Inspired Speech Envelope Extraction Methods for Improved EEG-Based Auditory Attention Detection in a Cocktail Party Scenario. IEEE Transactions on Neural Systems and Rehabilitation Engineering. 2017 May;25(5):402–12.

[5] Vandecappelle S, Deckers L, Das N, Ansari AH, Bertrand A, Francart T. EEG-based Detection of the Locus of Auditory Attention with Convolutional Neural Networks. eLife. 2021 Apr;10:e56481.

[6] Geirnaert S, Francart T, Bertrand A. Fast EEG-Based Decoding Of The Directional Focus Of Auditory Attention Using Common Spatial Patterns. IEEE Transactions on Biomedical Engineering. 2021 May;68(5):1557–68.

[7] Kong YY, Mullangi A, Ding N. Differential Modulation of Auditory Responses to Attended and Unattended Speech in Different Listening Conditions. Hearing Research. 2014 Oct;316:73–81.

[8] Vanthornhout J, Decruy L, Francart T. Effect of Task and Attention on Neural Tracking of Speech. Frontiers in Neuroscience. 2019;13:977.

[9] Lesenfants D, Francart T. The Interplay of Top-down Focal Attention and the Cortical Tracking of Speech. Scientific Reports. 2020 Dec;10(1):6922.

[10] Ding N, Simon JZ. Emergence of Neural Encoding of Auditory Objects While Listening to Competing Speakers. Proceedings of the National Academy of Sciences. 2012 Jul;109(29):11854–9.

[11] Zion Golumbic EM, Ding N, Bickel S, Lakatos P, Schevon CA, McKhann GM, et al. Mechanisms Underlying Selective Neuronal Tracking of Attended Speech at a ‘Cocktail Party’. Neuron. 2013 Mar;77(5):980–91.

[12] Puvvada KC, Simon JZ. Cortical Representations of Speech in a Multitalker Auditory Scene. Journal of Neuroscience. 2017 Sep;37(38):9189–96.

[13] Dougherty ER. Random Processes for Image Signal Processing. Bellingham: Wiley-IEEE Press; 1998.

[14] Viertiö-Oja H, Maja V, Särkelä M, Talja P, Tenkanen N, Tolvanen-Laakso H, et al. Description of the Entropy™ Algorithm as Applied in the Datex-Ohmeda S/5™ Entropy Module. Acta Anaesthesiologica Scandinavica. 2004;48(2):154–61.

[15] Rezek IA, Roberts SJ. Stochastic Complexity Measures for Physiological Signal Analysis. IEEE Transactions on Biomedical Engineering. 1998;45(9):1186–91.

[16] Lesenfants D, Habbal D, Chatelle C, Soddu A, Laureys S, Noirhomme Q. Toward an Attention-Based Diagnostic Tool for Patients With Locked-in Syndrome. Clinical EEG and Neuroscience. 2018 Mar;49(2):122–35.

[17] Inouye T, Shinosaki K, Sakamoto H, Toi S, Ukai S, Iyama A, et al. Quantification of EEG Irregularity by Use of the Entropy of the Power Spectrum. Electroencephalography and Clinical Neurophysiology. 1991 Sep;79(3):204–10.

[18] Belyavin A, Wright NA. Changes in Electrical Activity of the Brain with Vigilance. Electroencephalography and Clinical Neurophysiology. 1987 Feb;66(2):137–44.

[19] Geirnaert S, Francart T, Bertrand A. Unsupervised Self-Adaptive Auditory Attention Decoding. IEEE journal of biomedical and health informatics. 2021 Oct;25(10):3955–66.

[20] de Winter JCF, Gosling SD, Potter J. Comparing the Pearson and Spearman Correlation Coefficients across Distributions and Sample Sizes: A Tutorial Using Simulations and Empirical Data. Psychological Methods. 2016 Sep;21(3):273–90.

[21] Das N, Francart T, Bertrand A. Auditory Attention Detection Dataset KULeuven (1.0.0) [Data set]; 2019. https://zenodo.org/record/3377911.

[22] Mobile EEG for Neuroscience Reseach – Mbt | mBrainTrain; 2023. Available from: https://mbraintrain.com/.

[23] Lindgren J. Converting. Ov Files to Matlab; 2015. Available from: http://openvibe.inria.fr/converting-ov-files-to-matlab/.

[24] deBuren. Radioboeken voor kinderen; 2007. Available from: https://soundcloud.com/deburen-eu/sets/radioboeken-voor-kinderen.

[25] Dahl R. Alle verhalen. Amsterdam: Meulenhof; 2013.

[26] Patterson RD, Allerhand MH, Giguère C. Time-domain Modeling of Peripheral Auditory Processing: A Modular Architecture and a Software Platform. The Journal of the Acoustical Society of America. 1995 Oct;98(4):1890–4.

[27] Søndergaard P, Majdak P. The Auditory Modeling Toolbox. In: The Technology of Binaural Listening, Modern Acoustics and Signal Processing. Berlin: Springer; 2013. p. 33–56.

[28] Somers B, Francart T, Bertrand A. GitHub repository: MWF Toolbox for EEG Artifact Removal; 2023. Available from: https://github.com/exporl/mwf-artifact-removal.

[29] Somers B, Francart T, Bertrand A. A Generic EEG Artifact Removal Algorithm Based on the Multi-Channel Wiener Filter. Journal of Neural Engineering. 2018 Jun;15(3):036007.

[30] Delorme A, Sejnowski T, Makeig S. Enhanced Detection of Artifacts in EEG Data Using Higher-Order Statistics and Independent Component Analysis. NeuroImage. 2007;34(4):1443–9.

[31] Ledoit O, Wolf M. A Well-Conditioned Estimator for Large-Dimensional Covariance Matrices. Journal of Multivariate Analysis. 2004 Feb;88(2):365–411.

[32] Ledoit O, Wolf M. Honey, I Shrunk the Sample Covariance Matrix; 2014. Available from: http://ledoit.net/honey_abstract.htm.

[33] Babadi B, Brown EN. A Review of Multitaper Spectral Analysis. IEEE Transactions on Biomedical Engineering. 2014 May;61(5):1555–64.

[34] Hastie T, Tibshirani R, Friedman J. The Elements of Statistical Learning. 2nd ed. Springer Series in Statistics. New York: Springer; 2009.

[35] Wilcoxon F. Individual Comparisons by Ranking Methods. Biometrics Bulletin. 1945;1(6):80–3.

[36] Benjamini Y, Hochberg Y. Controlling the False Discovery Rate: A Practical and Powerful Approach to Multiple Testing. Journal of the Royal Statistical Society Series B (Methodological). 1995;57(1):289–300.

[37] Student. The Probable Error of a Mean. Biometrika. 1908;6(1):1–25.

[38] Tune S, Wöstmann M, Obleser J. Probing the limits of alpha power lateralisation as a neural marker of selective attention in middle-aged and older listeners. European Journal of Neuroscience. 2018;48(7):2537–50.

[39] Rotaru I, Geirnaert S, Heintz N, de Ryck IV, Bertrand A, Francart T. EEG-based decoding of the spatial focus of auditory attention in a multi-talker audiovisual experiment using Common Spatial Patterns. bioRxiv. 2023. Available from: https://www.biorxiv.org/content/early/2023/07/15/2023.07.13.548824.

[40] Puffay C, Accou B, Bollens L, Monesi MJ, Vanthornhout J, hamme HV, et al. Relating EEG to Continuous Speech Using Deep Neural Networks: A Review. Journal of Neural Engineering. 2023 Aug;20(4):041003.

[41] de Cheveigné A, Wong DDE, Di Liberto GM, Hjortkjær J, Slaney M, Lalor E. Decoding the Auditory Brain with Canonical Component Analysis. NeuroImage. 2018 May;172:206–16.

[42] de Cheveigné A, Slaney M, Fuglsang SA, Hjortkjaer J. Auditory Stimulus-Response Modeling with a Match-Mismatch Task. Journal of Neural Engineering. 2021 Aug;18(4):046040.

